## Supplemental Information for "MetaTiME: Meta-components of the Tumor Immune Microenvironment"

### Supplementary Figures

#### Fig.S1.

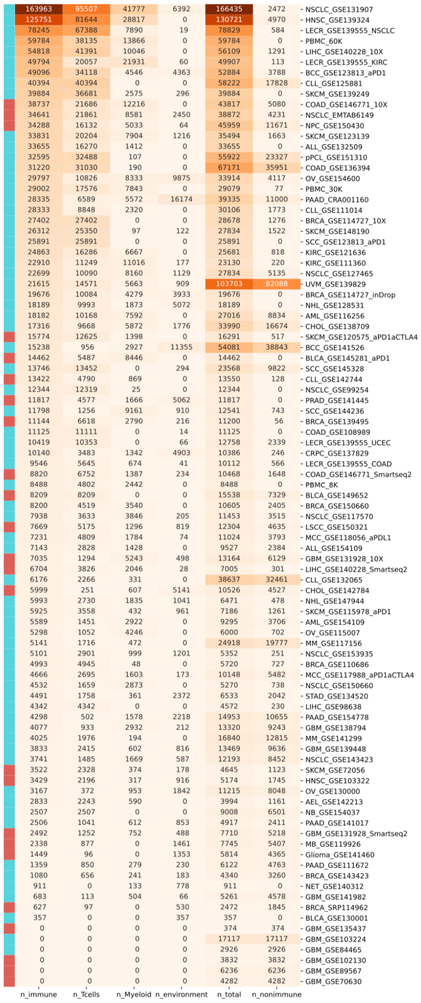


**Fig.S1. Cell statistics of public tumor scRNA-seq collection.** We utilized the curated cell type annotation from TISCH to summarize the number of cells in each category. MetaTiME includes immune cells and tumor microenvironment cells from each dataset, and excludes the malignant cells with low reproducibility across datasets. Datasets with raw counts available are marked as blue while datasets with only continuous TPM available are marked as red.

**Fig.S2.**

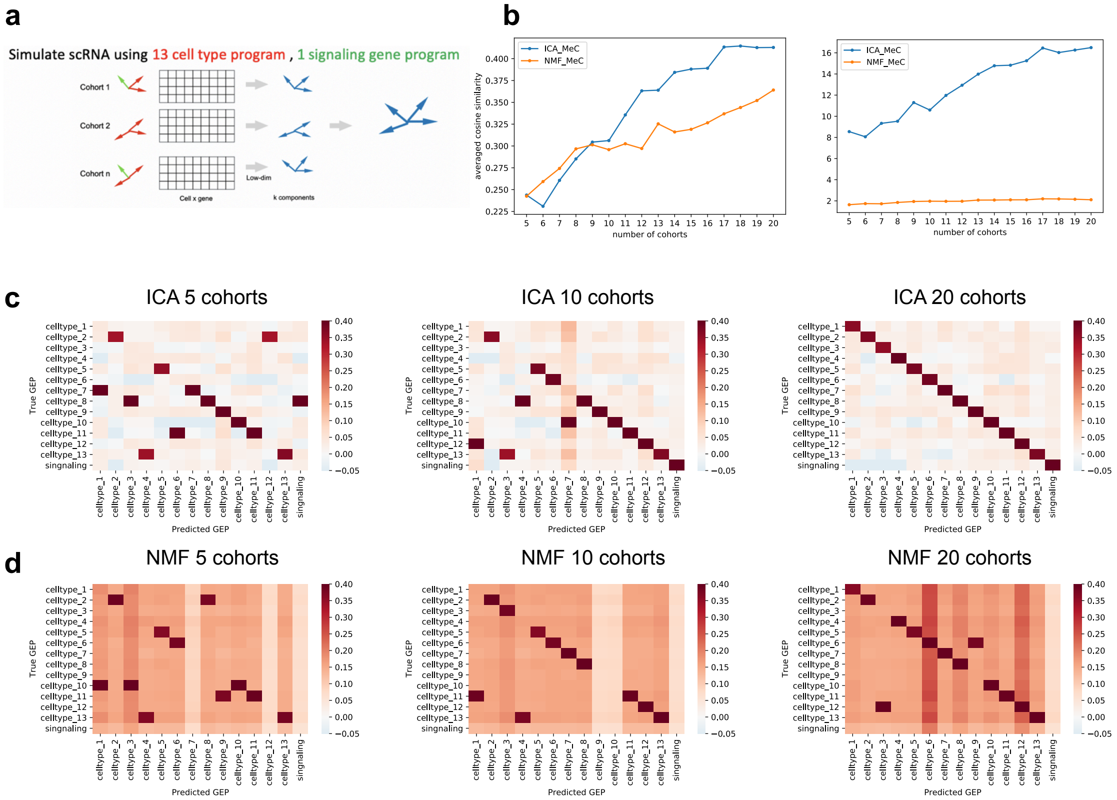


**Fig.S2. Simulating scRNA data with transcriptional gene programs.** Two low-dimensional reduction methods were benchmarked using simulated scRNA data **(a)** Simulated gene expression program includes 13 cell type-specific gene programs and one pan-cell type gene program, mimicking common signaling pathways in single-cell RNA-seq data. The MetaTiME method was applied on the simulated cohorts to extract meta-components (MeCs) which were later compared with the pre-defined gene programs using correlation. **(b)** Numbers of cohorts were tested to assess the robustness of meta-components. The similarity between the recovered gene programs and the true gene programs (GEP) increases with larger cohort numbers. The average similarity score of ICA-derived MeCs is higher than NMF-derived MeCs. **(c)** The pairwise similarity between true GEP and predicted GEP using ICA for 5, 10, 20 cohorts. **(d)** The pairwise similarity between true GEP and predicted GEP using NMF for 5, 10, 20 cohorts.

**Fig.S3.**

a

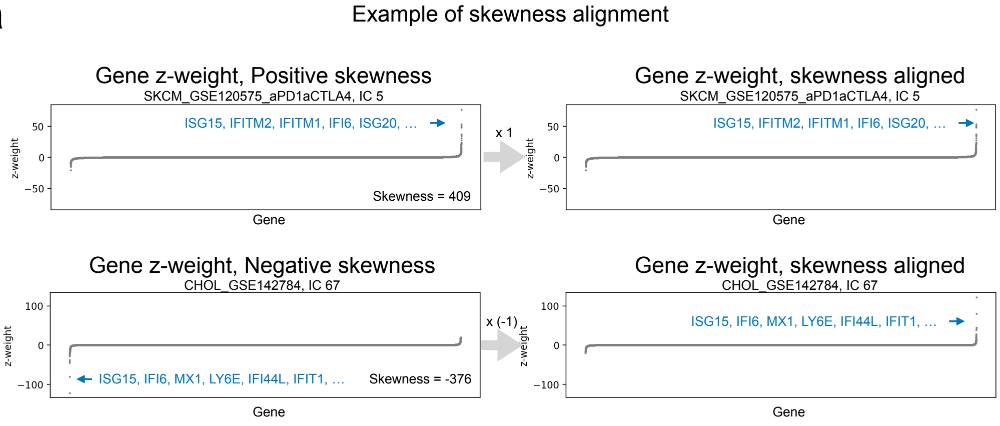


b

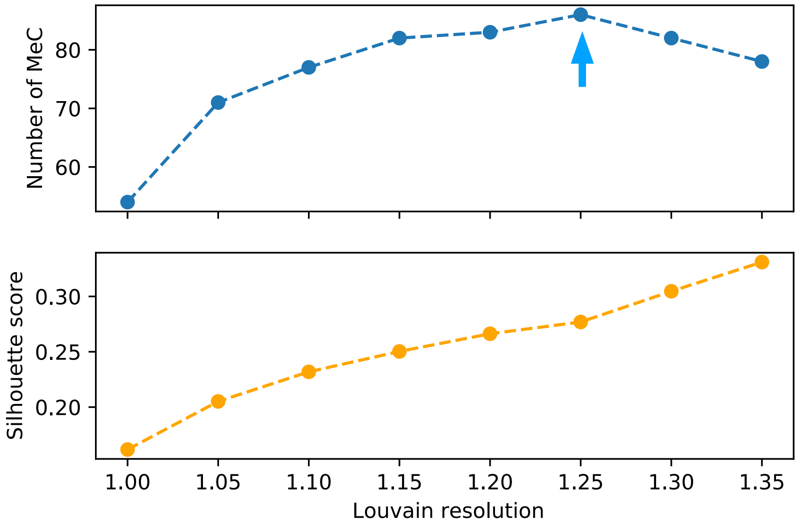


**Fig.S3. Meta-component skewness and parameter tuning. (a)** Genes most highly associated with a meta-component (MeC) tend to be biased to either significantly positive or significantly negative z-weights since the signs of the independent components are arbitrary. Examples of skewness of two MeCs, one with positive skewness (top row), the other with negative skewness (bottom row). MeCs with negative skewness are flipped by multiplying the gene weights by -1. **(b)** To determine the number of MeCs, we tested a range of resolution parameters in the community-detection based Louvain clustering algorithm. The Silhouette score increases as resolution increases. However, higher Louvain clustering resolution results in MeC clusters that are filtered as insufficiently reproducible across cohorts, which results in an overall decrease in the MeC number. The resolution parameter was thus selected to be 1.25, leading to 86 MeCs that optimize the tradeoff between cluster separation and reproducibility.

Fig.S4.

a

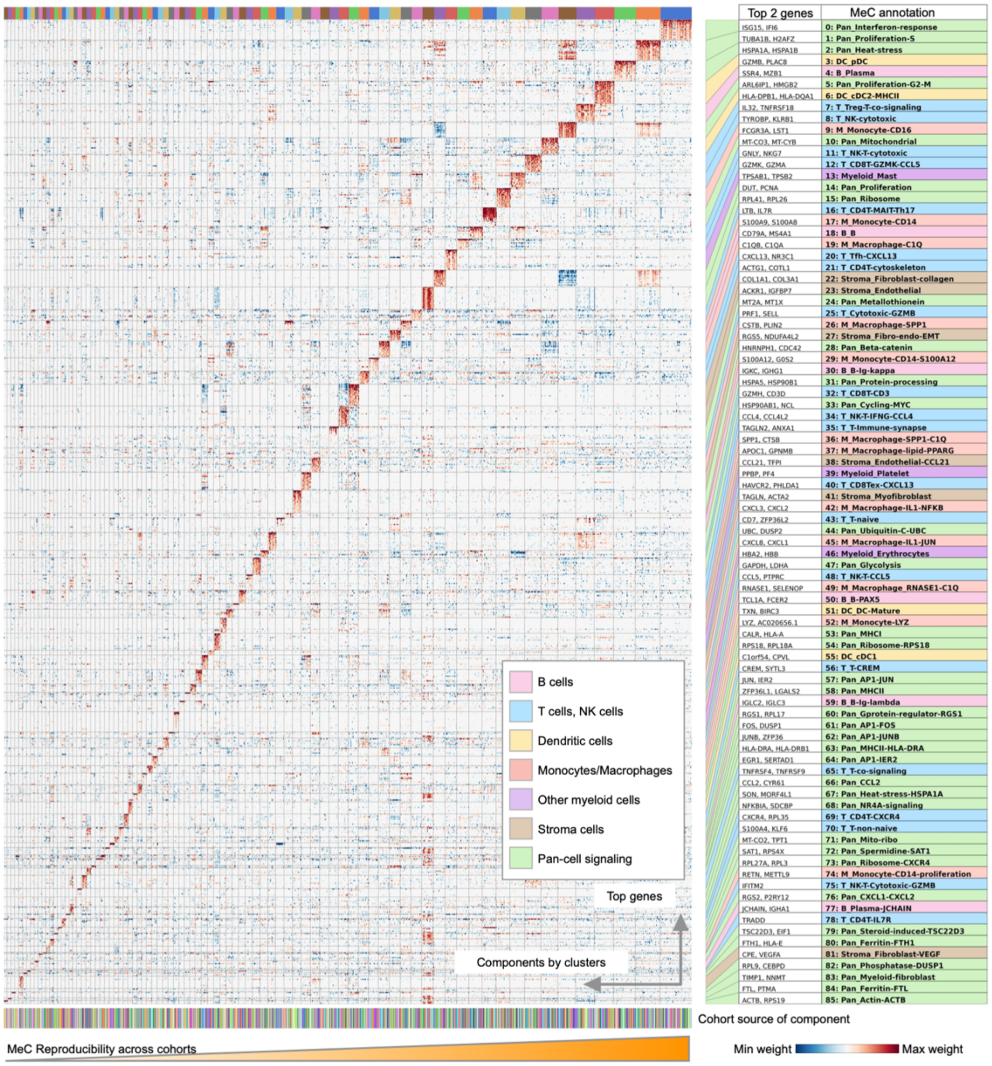

Top 2 genes
MeC annotation
ISG15, IFI6 | 0: Pan_Interferon-response
TUBA1B, H2AFZ | 1: Pan_Proliferation-S
HSPA1A, HSPA1B | 2: Pan_Heat-stress
GZMB, PLAC8 | 3: DC_pDC
SSR4, MZB1 | 4: B_Plasma
ARL6IP1, HMGB2 | 5: Pan_Proliferation-G2-M
HLA-DPB1, HLA-DQA1 | 6: DC_cDC2-MHCII
IL32, TNFRSF18 | 7: T_Treg-T-co-signaling
TYROBP, KLRB1 | 8: T_NK-cytotoxic
FCGR3A, LST1 | 9: M_Monocyte-CD16
MT-CO3, MT-CYB | 10: Pan_Mitochondrial
GNLY, NKG7 | 11: T_NK-T-cytotoxic
GZMK, GZMA | 12: T_CD8T-GZMK-CCL5
TPSAB1, TPSB2 | 13: Myeloid_Mast
DUT, PCNA | 14: Pan_Proliferation
RPL41, RPL26 | 15: Pan_Ribosome
LTB, IL7R | 16: T_CD4T-MAIT-Th17
S100A9, S100A8 | 17: M_Monocyte-CD14
CD79A, MS4A1 | 18: B_B
C1QB, C1QA | 19: M_Macrophage-C1Q
CXCL13, NR3C1 | 20: T_Tfh-CXCL13
ACTG1, COTL1 | 21: T_CD4T-cytoskeleton
COL1A1, COL3A1 | 22: Stroma_Fibroblast-collagen
ACKR1, IGFBP7 | 23: Stroma_Endothelial
MT2A, MT1X | 24: Pan_Metallothionein
PRF1, SELL | 25: T_Cytotoxic-GZMB
CSTB, PLIN2 | 26: M_Macrophage-SPP1
RGS5, NDUFA4L2 | 27: Stroma_Fibro-endo-EMT
HNRNPH1, CDC42 | 28: Pan_Beta-catenin
S100A12, G0S2 | 29: M_Monocyte-CD14-S100A12
IGKC, IGHG1 | 30: B_B-Ig-kappa
HSPA5, HSP90B1 | 31: Pan_Protein-processing
GZMH, CD3D | 32: T_CD8T-CD3
HSP90AB1, NCL | 33: Pan_Cycling-MYC
CCL4, CCL4L2 | 34: T_NK-T-IFNG-CCL4
TAGLN2, ANXA1 | 35: T_T-Immune-synapse
SPP1, CTSB | 36: M_Macrophage-SPP1-C1Q
APOC1, GPNMB | 37: M_Macrophage-lipid-PPARG
CCL21, TFPI | 38: Stroma_Endothelial-CCL21
PPBP, PF4 | 39: Myeloid_Platelet
HAVCR2, PHLDA1 | 40: T_CD8Tex-CXCL13
TAGLN, ACTA2 | 41: Stroma_Myofibroblast
CXCL3, CXCL2 | 42: M_Macrophage-IL1-NFKB
CD7, ZFP36L2 | 43: T_T-naive
UBC, DUSP2 | 44: Pan_Ubiquitin-C-UBC
CXCL8, CXCL1 | 45: M_Macrophage-IL1-JUN
HBA2, HBB | 46: Myeloid_Erythrocytes
GAPDH, LDHA | 47: Pan_Glycolysis
CCL5, PTPRC | 48: T_NK-T-CCL5
RNASE1, SELENOP | 49: M_Macrophage_RNASE1-C1Q
TCL1A, FCER2 | 50: B_B-PAX5
TXN, BIRC3 | 51: DC_DC-Mature
LYZ, AC020656.1 | 52: M_Monocyte-LYZ
CALR, HLA-A | 53: Pan_MHCI
RPS18, RPL18A | 54: Pan_Ribosome-RPS18
C1orf54, CPVL | 55: DC_cDC1
CREM, SYTL3 | 56: T_T-CREM
JUN, IER2 | 57: Pan_AP1-JUN
ZFP36L1, LGALS2 | 58: Pan_MHCII
IGLC2, IGLC3 | 59: B_B-Ig-lambda
RGS1, RPL17 | 60: Pan_Gprotein-regulator-RGS1
FOS, DUSP1 | 61: Pan_AP1-FOS
JUNB, ZFP36 | 62: Pan_AP1-JUNB
HLA-DRA, HLA-DRB1 | 63: Pan_MHCII-HLA-DRA
EGR1, SERTAD1 | 64: Pan_AP1-IER2
TNFRSF4, TNFRSF9 | 65: T_T-co-signaling
CCL2, CYR61 | 66: Pan_CCL2
SON, MORF4L1 | 67: Pan_Heat-stress-HSPA1A
NFKBIA, SDCBP | 68: Pan_NR4A-signaling
CXCR4, RPL35 | 69: T_CD4T-CXCR4
S100A4, KLF6 | 70: T_T-non-naive
MT-CO2, TPT1 | 71: Pan_Mito-ribo
SAT1, RPS4X | 72: Pan_Spermidine-SAT1
RPL27A, RPL3 | 73: Pan_Ribosome-CXCR4
RETN, METTL9 | 74: M_Monocyte-CD14-proliferation
IFITM2 | 75: T_NK-T-Cytotoxic-GZMB
RGS2, P2RY12 | 76: Pan_CXCL1-CXCL2
JCHAIN, IGHA1 | 77: B_Plasma-JCHAIN
TRADD | 78: T_CD4T-IL7R
TSC22D3, EIF1 | 79: Pan_Steroid-induced-TSC22D3
FTH1, HLA-E | 80: Pan_Ferritin-FTH1
CPE, VEGFA | 81: Stroma_Fibroblast-VEGF
RPL9, CEBPD | 82: Pan_Phosphatase-DUSP1
TIMP1, NNMT | 83: Pan_Myeloid-fibroblast
FTL, PTMA | 84: Pan_Ferritin-FTL
ACTB, RPS19 | 85: Pan_Actin-ACTB
B cells
T cells, NK cells
Dendritic cells
Monocytes/Macrophages
Other myeloid cells
Stroma cells
Pan-cell signaling
Top genes
Components by clusters
Cohort source of component
MeC Reproducibility across cohorts
Min weight
Max weight


b

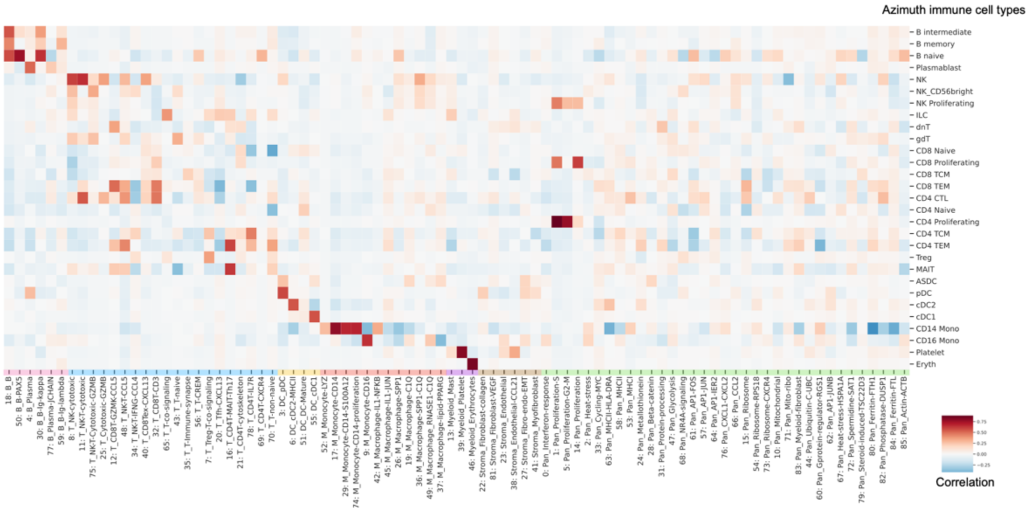

Azimuth immune cell types
B intermediate
B memory
B naive
Plasmablast
NK
NK_CD56bright
NK Proliferating
ILC
dnT
gdT
CD8 Naive
CD8 Proliferating
CD8 TCM
CD8 TEM
CD4 CTL
CD4 Naive
CD4 Proliferating
CD4 TCM
CD4 TEM
Treg
MAIT
ASDC
pDC
cDC2
cDC1
CD14 Mono
CD16 Mono
Platelet
Eryth
Correlation

**Fig.S4. Meta-components with annotation and comparison with known immune cell types. (a)** All independent components clustered in MeCs are plotted using the normalized MeC gene z-weights, showing top featured genes in each MeC. Each row represents a gene, and each column represents an independent component. Components are ordered by MeC cluster assignment, and genes are ranked using MeC z-weights and deduplicated. The MeC annotation of each of the 86 MeCs are shown on the right with the top 2 genes marked. **(b)** Correlation between all functional MeCs and Azimuth by converting the marker list defined for immune cell types and subtypes to a list of weight 1.

**Fig.S5.**

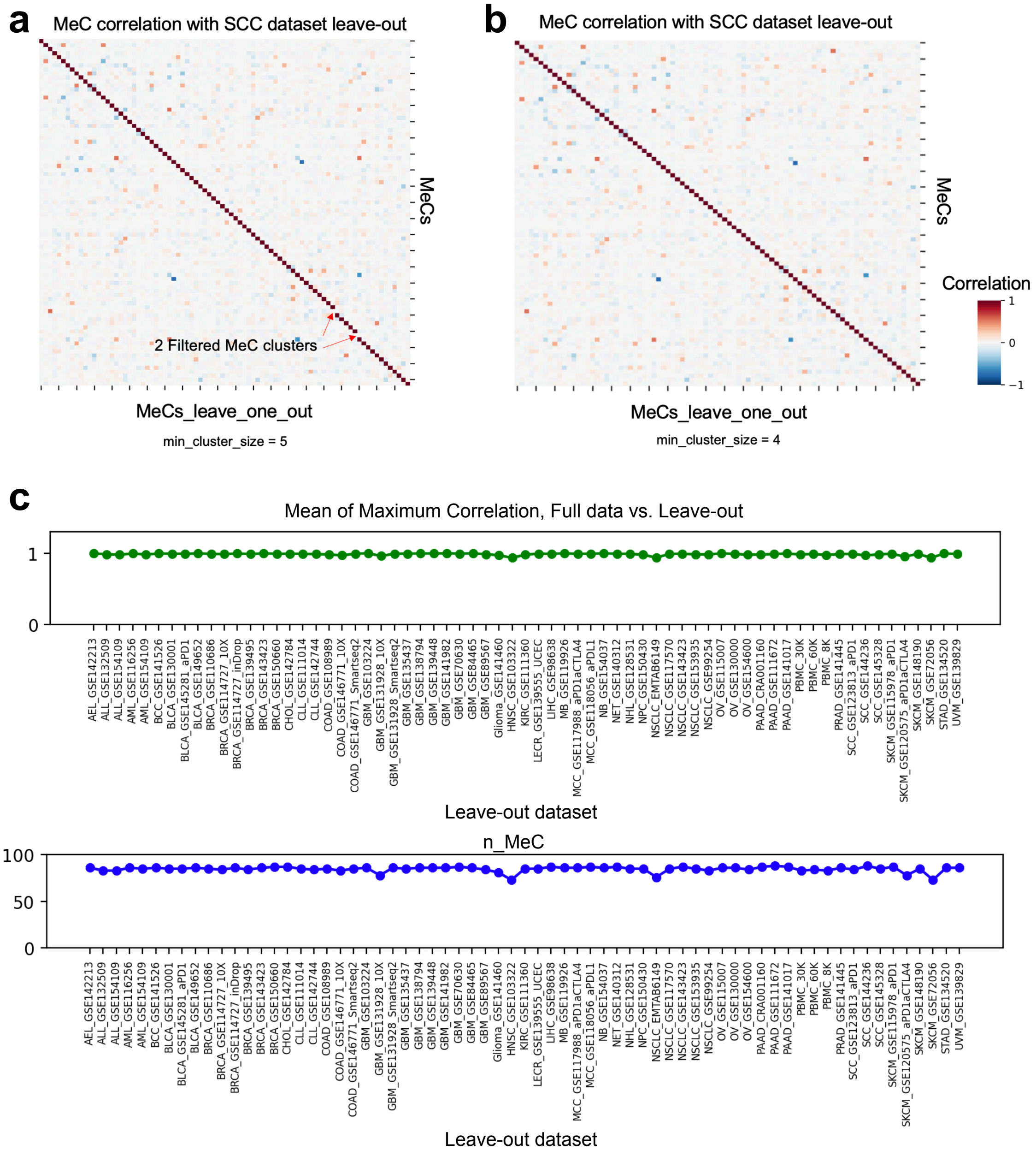


**Fig.S5. Leave-one-out MeC training. (a)** Correlation between MeCs called from the full set of datasets (row-wise) and the MeCs called leaving the SCC dataset out of MetaTiME training (column-wise), with the same cluster calling resolution parameter, and the minimum IC number in one cluster set to be 5 **(b)** Correlation between MeCs called from the full set (row-wise) and the MeCs called leaving the SCC dataset out (column-wise) when changing the minimum IC number in one cluster to be 4. **(c)** Leave-one-out MeCs compared to full-set MeCs, using the mean of row-wise and column-wise maximum correlation (top panel, mean max correlation=0.988), and number of MeCs (bottom, mean leave-one-dataset-out MeC number=84.64 compared to full-set MeC 86).

**Fig.S6.**

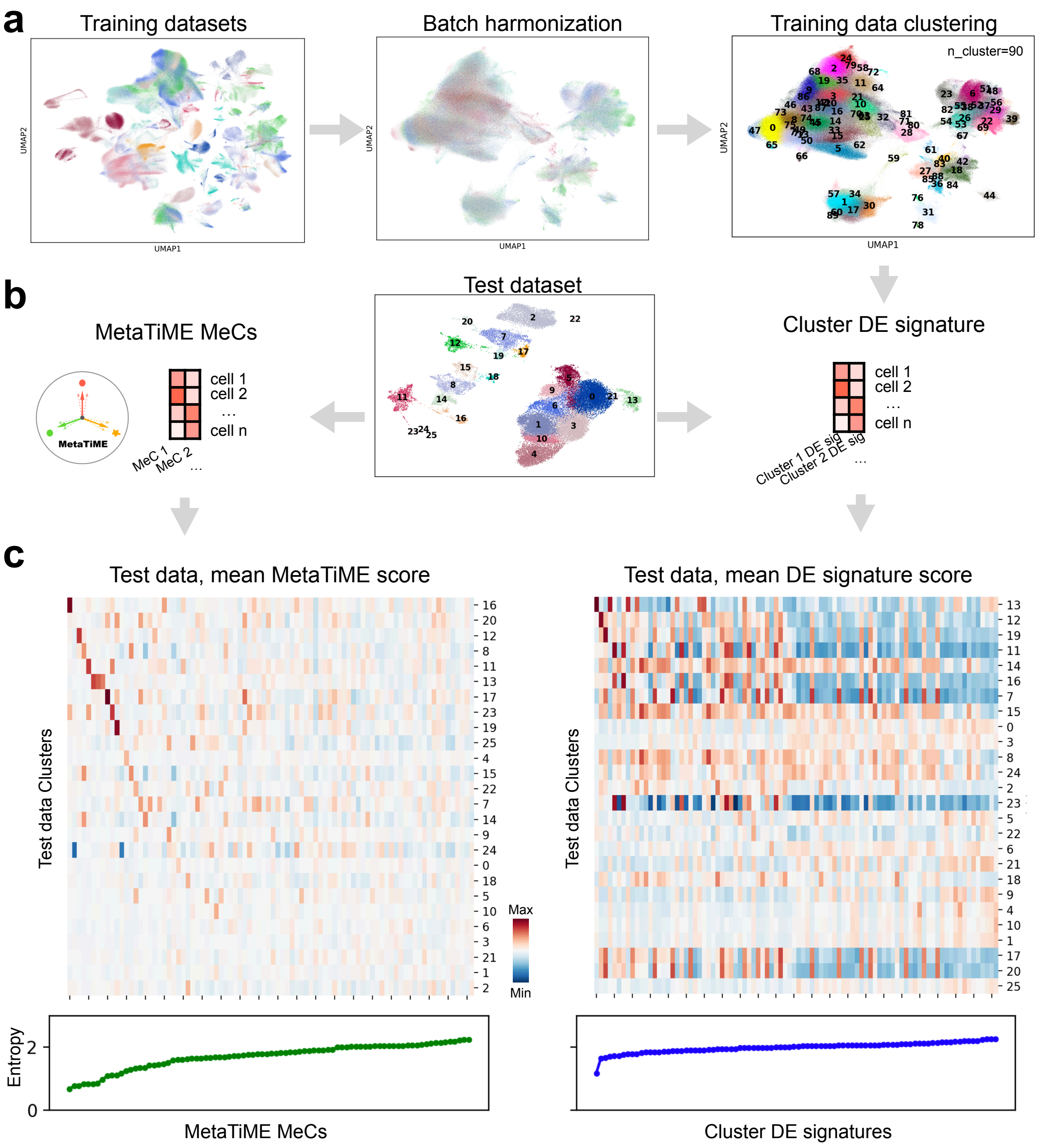


**Fig.S6. Comparison of MetaTiME MeCs to signatures generated by clustering and differential gene expression analysis. (a)** Direct integration of cells allows for inclusion of up to 21 datasets with the largest numbers of TME cells. Cells are harmonized across datasets as batches, clustered, and differential gene expression analysis performed to generate Cluster differential expression (DE) signatures. **(b)** Cluster DE signatures and MeC signatures are mapped to the test dataset where cells are also clustered. **(c)** The per-cluster mean scores of both signatures are plotted in heatmaps to assess whether the signatures are specific to test cell clusters or uniformly distributed, with entropy calculated to reflect the test cluster distribution of scores from MeC(left) and Cluster DE signatures (right).

**Fig.S7.**

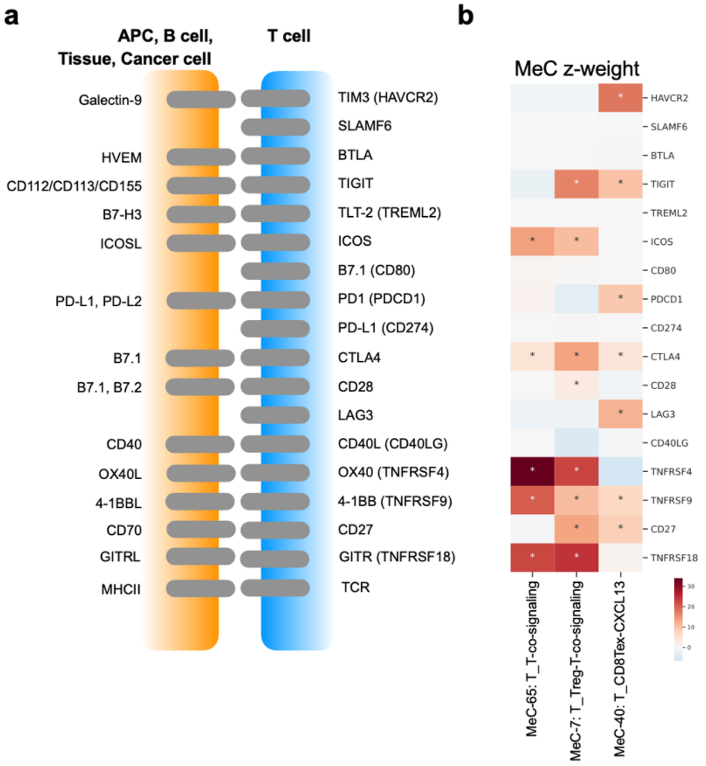


**c**

| MeC-65: T_T-co-signaling | z-weight | rank |
|---|---|---|
| TNFRSF4 | 33.7 | 0 |
| TNFRSF18 | 22.6 | 1 |
| TNFRSF9 | 20.4 | 2 |
| ICOS | 14.0 | 9 |
| CTLA4 | 4.5 | 108 |
| TNFRSF8 | 4.3 | 117 |

| MeC-7: T_Treg-T-co-signaling | z-weight | rank |
|---|---|---|
| HAVCR2 | 18.1 | 5 |
| LAG3 | 11.8 | 17 |
| TIGIT | 9.8 | 28 |
| PDCD1 | 9.1 | 33 |
| CD27 | 8.0 | 42 |
| TNFRSF9 | 7.2 | 48 |
| CTLA4 | 4.2 | 100 |

| MeC-40: T_CD8Tex-CXCL13 | z-weight | rank |
|---|---|---|
| TNFRSF18 | 24.5 | 1 |
| TNFRSF4 | 22.7 | 2 |
| TIGIT | 16.8 | 11 |
| CD27 | 13.4 | 14 |
| CTLA4 | 13.2 | 15 |
| ICOS | 10.6 | 21 |
| TNFRSF9 | 10.3 | 24 |
| CD28 | 3.2 | 127 |

**Fig.S7. MetaTiME meta-components reveal co-signaling pathway genes in T cells. (a)** Illustration of genes encoding membrane receptors on the T cell on the T cell co-stimulating and co-inhibitory pathways, with knowledge of interacting ligands from antigen presenting cells (APC), B cells, tissue cells or cancer cells. **(b)** Heatmap of meta-component z-weight of T co-signaling genes in three related T cell MeCs. Genes with significant contributions to the MeC are marked with a star. **(c)** Z-weight and ranking of T co-signaling genes in three related T cell MeCs.

**Fig.S8.**

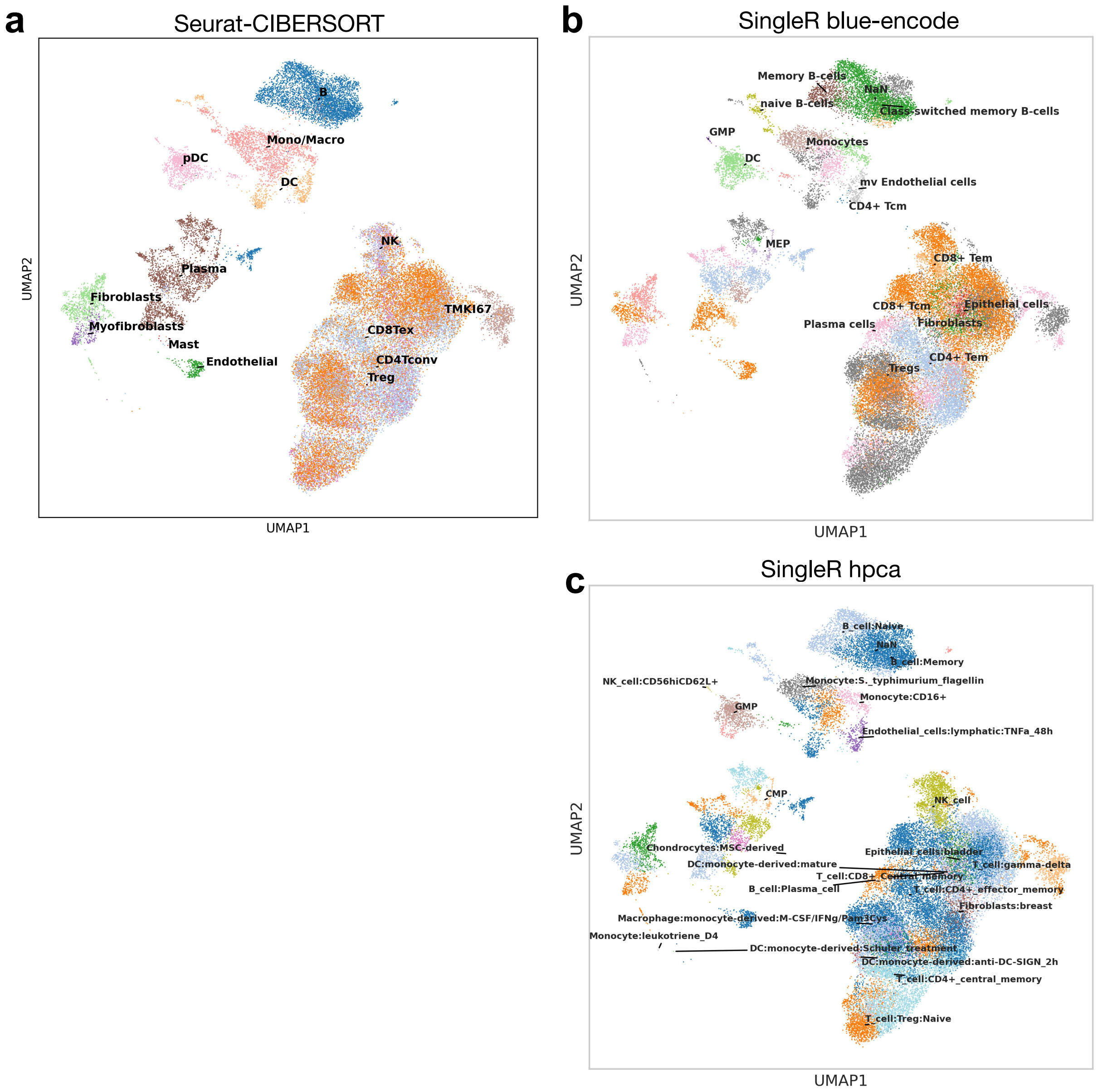


**Fig.S8. Automatic annotation using existing methods and marker panels. (a).** Seurat annotated results when using the CIBERSORT panel, as implemented in MAESTRO. **(b).** SingleR annotated results using the immune related panel Blueprint ENCODE (blue-encode). **(c).** SingleR annotated results using the immune related panel Human Primary Cell Atlas (hpca).

### Fig.S9.

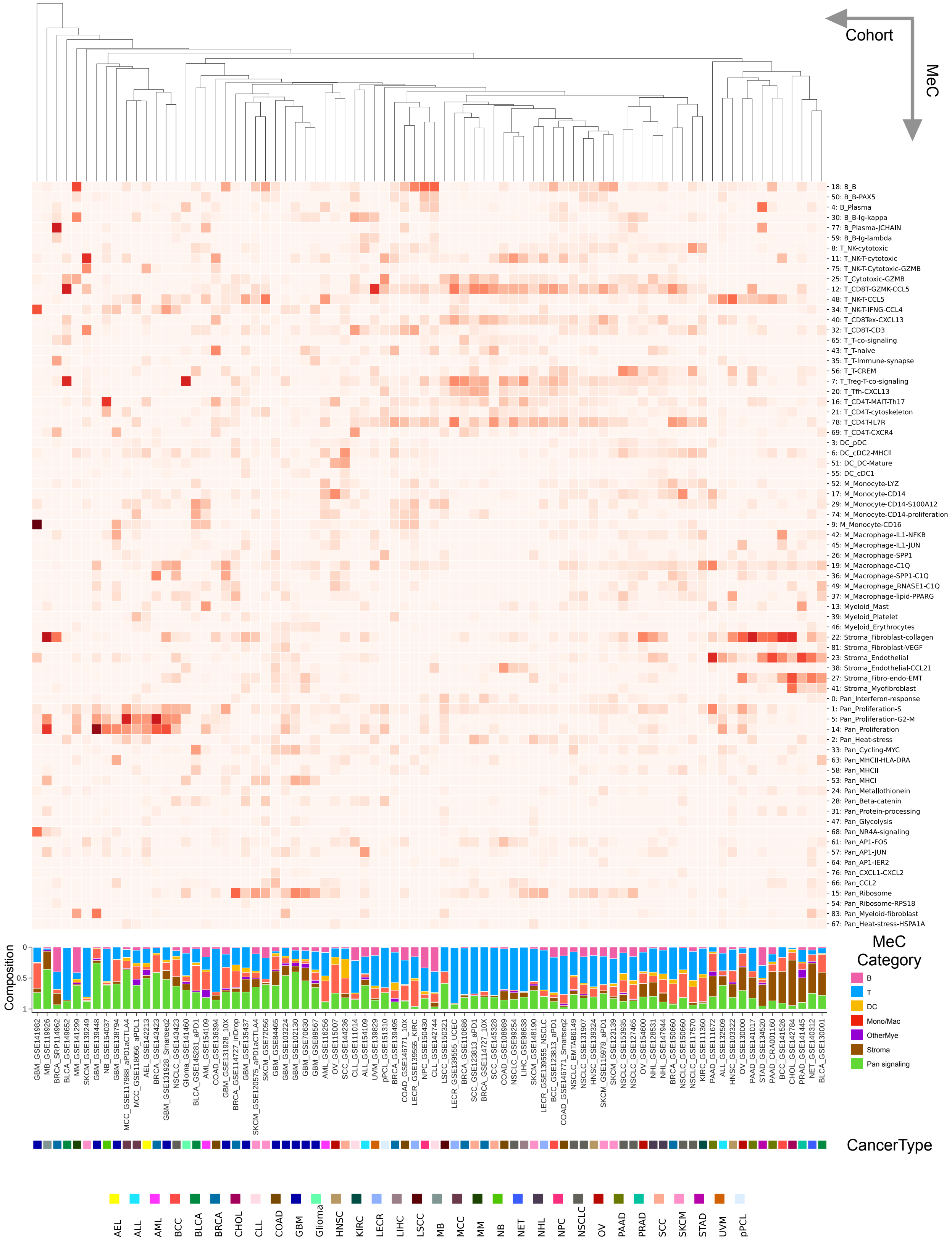
Cohort
MeC
18: B_B
50: B_B-PAX5
4: B_Plasma
30: B_B-Ig-kappa
77: B_Plasma-JCHAIN
59: B_B-Ig-lambda
8: T_NK-cytotoxic
11: T_NK-T-cytotoxic
75: T_NK-T-Cytotoxic-GZMB
25: T_Cytotoxic-GZMB
12: T_CD8T-GZMK-CCL5
48: T_NK-T-CCL5
34: T_NK-T-IFNG-CCL4
40: T_CD8Tex-CXCL13
32: T_CD8T-CD3
65: T_T-co-signaling
43: T_T-naive
35: T_T-Immune-synapse
56: T_T-CREM
7: T_Treg-T-co-signaling
20: T_Tfh-CXCL13
16: T_CD4T-MAIT-Th17
21: T_CD4T-cytoskeleton
78: T_CD4T-IL7R
69: T_CD4T-CXCR4
3: DC_pDC
6: DC_cDC2-MHCII
51: DC_DC-Mature
55: DC_cDC1
52: M_Monocyte-LYZ
17: M_Monocyte-CD14
29: M_Monocyte-CD14-S100A12
74: M_Monocyte-CD14-proliferation
9: M_Monocyte-CD16
42: M_Macrophage-IL1-NFKB
45: M_Macrophage-IL1-JUN
26: M_Macrophage-SPP1
19: M_Macrophage-C1Q
36: M_Macrophage-SPP1-C1Q
49: M_Macrophage_RNASE1-C1Q
37: M_Macrophage-lipid-PPARG
13: Myeloid_Mast
39: Myeloid_Platelet
46: Myeloid_Erythrocytes
22: Stroma_Fibroblast-collagen
81: Stroma_Fibroblast-VEGF
23: Stroma_Endothelial
38: Stroma_Endothelial-CCL21
27: Stroma_Fibro-endo-EMT
41: Stroma_Myofibroblast
0: Pan_Interferon-response
1: Pan_Proliferation-S
5: Pan_Proliferation-G2-M
14: Pan_Proliferation
2: Pan_Heat-stress
33: Pan_Cycling-MYC
63: Pan_MHCII-HLA-DRA
58: Pan_MHCII
53: Pan_MHCI
24: Pan_Metallothionein
28: Pan_Beta-catenin
31: Pan_Protein-processing
47: Pan_Glycolysis
68: Pan_NR4A-signaling
61: Pan_AP1-FOS
57: Pan_AP1-JUN
64: Pan_AP1-IER2
76: Pan_CXCL1-CXCL2
66: Pan_CCL2
15: Pan_Ribosome
54: Pan_Ribosome-RPS18
83: Pan_Myeloid-fibroblast
67: Pan_Heat-stress-HSPA1A
Composition
0
0.5
1
MeC Category
B
T
DC
Mono/Mac
OtherMye
Stroma
Pan signaling
GBM_GSE141982
MB_GSE119926
BRCA_SRP114962
BLCA_GSE149652
MM_GSE141299
SKCM_GSE139249
GBM_GSE139448
NB_GSE154037
GBM_GSE138794
MCC_GSE117988_aPD1aCTLA4
MCC_GSE118056_aPDL1
AEL_GSE142213
BRCA_GSE143423
GBM_GSE131928_Smartseq2
NSCLC_GSE143423
Glioma_GSE141460
BLCA_GSE145281_aPDL1
AML_GSE154109
COAD_GSE136394
GBM_GSE131928_10X
BRCA_GSE114727_inDrop
GBM_GSE135437
SKCM_GSE120575_aPD1aCTLA4
SKCM_GSE72056
GBM_GSE84465
GBM_GSE103224
GBM_GSE102130
GBM_GSE70630
GBM_GSE89567
AML_GSE116256
OV_GSE115007
SCC_GSE144236
CLL_GSE111014
ALL_GSE154109
UVM_GSE139829
pPCL_GSE151310
BRCA_GSE139495
COAD_GSE146771_10X
LECR_GSE139555_KIRC
NPC_GSE150430
CLL_GSE125881
LSCC_GSE150321
LECR_GSE139555_UCEC
BRCA_GSE110686
SCC_GSE123813_aPD1
BRCA_GSE114727_10X
SCC_GSE145328
COAD_GSE108989
NSCLC_GSE99254
LIHC_GSE98638
SKCM_GSE148190
LECR_GSE139555_NSCLC
BCC_GSE123813_aPD1
COAD_GSE146771_Smartseq2
NSCLC_EMTAB6149
NSCLC_GSE131907
HNSC_GSE139324
SKCM_GSE115978_aPD1
SKCM_GSE123139
NSCLC_GSE153935
NSCLC_GSE127465
OV_GSE154600
NHL_GSE128531
NHL_GSE147944
BRCA_GSE150660
NSCLC_GSE150660
NSCLC_GSE117570
KIRC_GSE111360
PAAD_GSE111672
ALL_GSE132509
HNSC_GSE103322
OV_GSE130000
PAAD_GSE141017
STAD_GSE134520
PAAD_CRA001160
BCC_GSE141526
CHOL_GSE142784
PRAD_GSE141445
NET_GSE140312
BLCA_GSE130001
CancerType
AEL
ALL
AML
BCC
BLCA
BRCA
CHOL
CLL
COAD
GBM
Glioma
HNSC
KIRC
LECR
LIHC
LSCC
MB
MCC
MM
NB
NET
NHL
NPC
NSCLC
OV
PAAD
PRAD
SCC
SKCM
STAD
UVM
pPCL

**Fig.S9. Pan-cancer annotation of cell state composition in the full tumor scRNA datasets.** Top: heatmap showing cell proportions in each dataset. Bottom: bar plot showing cell state composition of tumor microenvironment for tumor scRNA dataset cell states. The proportion of cell states from the same MeC category are aggregated.

**Fig.S10.**

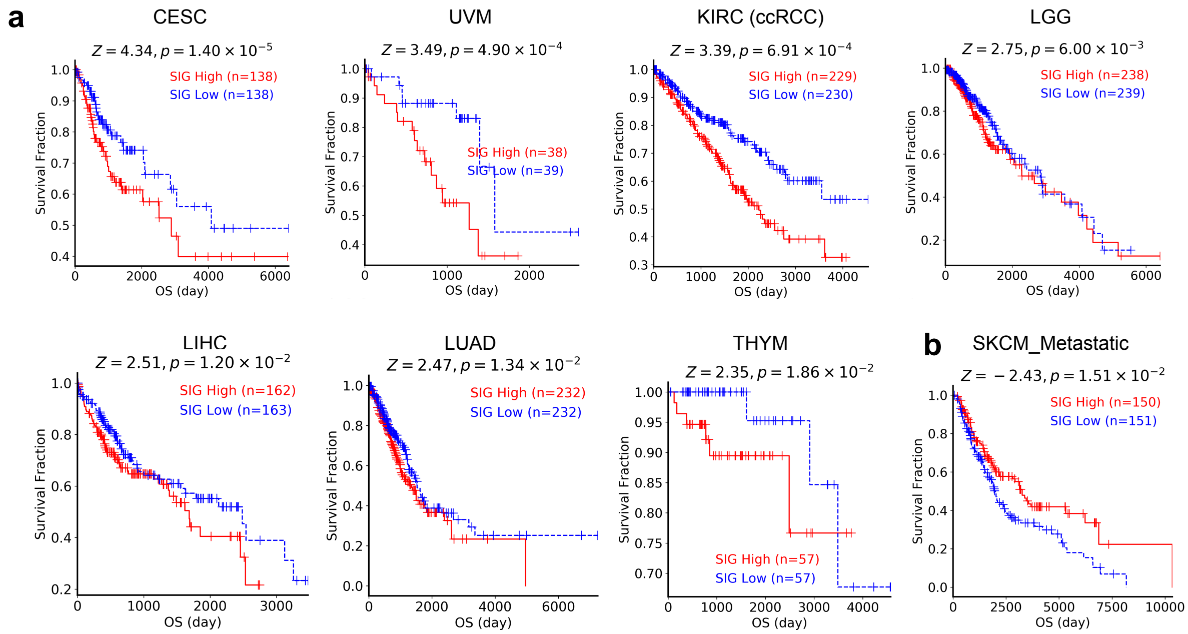


*SIG: signature of the MeC: 42: M_Macrophage-IL1-NFKB

**Fig.S10. TCGA survival prognosis of MeC representing IL1-activated macrophages.** Each sample in TCGA is evaluated using top genes in MeC-42 “M_Macrophage-IL1-NFKB”, and patients are separated into two groups based on expression level of the signature. Survival fractions of patients are shown for all cancer types with significant separation. **(a)** In seven cancer types, patients with higher signatures have worse prognosis. **(b)** In metastatic melanoma, patients with lower signatures have worse prognoses. CESC: cervical squamous cell carcinoma and endocervical adenocarcinoma. UVM: uveal melanoma. ccRCC: clear cell renal cell carcinoma, named as KIRC in TCGA. LGG: low grade glioma. LIHC: liver hepatocellular carcinoma. LUAD: lung adenocarcinoma. THYM: thymoma. SKCM: skin cutaneous melanoma.

**Fig.S11.**

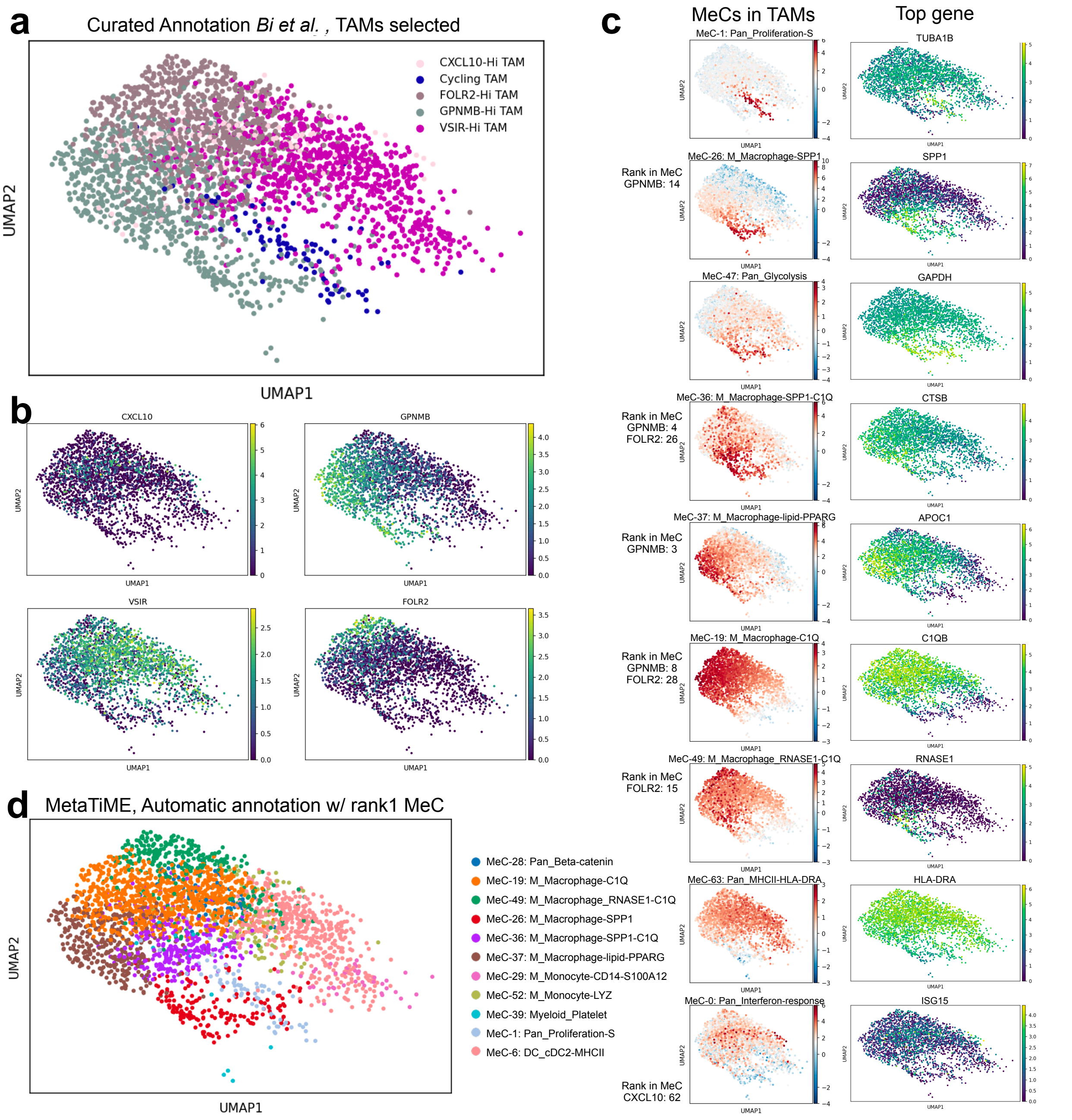


**Fig.S11. Comparison with myeloid markers in a previous kidney cancer study**. **(a)** Analysis was done on a kidney cohort with tumor associated macrophages. Cell types from the original study are shown. **(b)** Expression level of cell type markers used in the previous study. **(c)** Each row shows the scoring of a monocyte or macrophage related MetaTiME meta-component signature with expression of the marker gene top ranked by the meta-component. **(d)** A cluster-wise summary view of the most enriched MetaTiME meta-component. TAM: tumor associated macrophages.

**Fig.S12.**

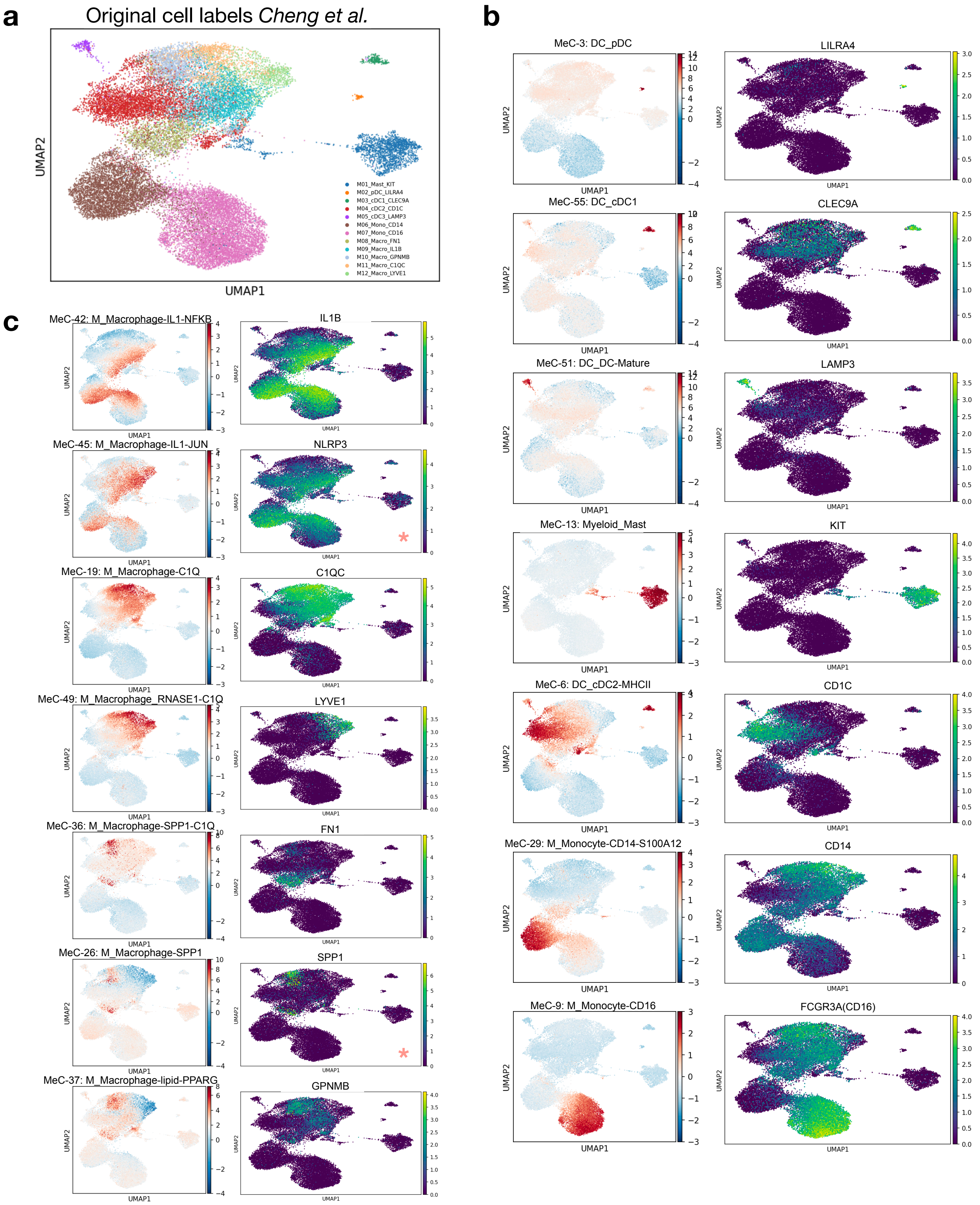
a
Original cell labels Cheng et al.
UMAP2
UMAP1
M01_Mast_KIT
M02_pDC_LILRA4
M03_cDC1_CLEC9A
M04_cDC2_CD1C
M05_cDC3_LAMP3
M06_Mono_CD14
M07_Mono_CD16
M08_Macro_FN1
M09_Macro_IL1B
M10_Macro_GPNMB
M11_Macro_C1QC
M12_Macro_LYVE1
b
MeC-3: DC_pDC
LILRA4
MeC-55: DC_cDC1
CLEC9A
MeC-51: DC_DC-Mature
LAMP3
MeC-13: Myeloid_Mast
KIT
MeC-6: DC_cDC2-MHCII
CD1C
MeC-29: M_Monocyte-CD14-S100A12
CD14
MeC-9: M_Monocyte-CD16
FCGR3A(CD16)
c
MeC-42: M_Macrophage-IL1-NFKB
IL1B
MeC-45: M_Macrophage-IL1-JUN
NLRP3
MeC-19: M_Macrophage-C1Q
C1QC
MeC-49: M_Macrophage_RNASE1-C1Q
LYVE1
MeC-36: M_Macrophage-SPP1-C1Q
FN1
MeC-26: M_Macrophage-SPP1
SPP1
MeC-37: M_Macrophage-lipid-PPARG
GPNMB
UMAP1
UMAP2

**Fig.S12. Comparison with myeloid types defined by clustering and selected markers in a previous myeloid-focused kidney cancer study. (a)** Analysis was done on a kidney myeloid cohort. Cell types from the original study are shown. **(b), (c)** Each row shows the scoring of a related MetaTiME myeloid meta-component signature (left) with expression of the marker gene from the original study (right). Genes with an orange star are marker genes from top of the MeC but not used in the original kidney cell types.

**Fig.S13.**

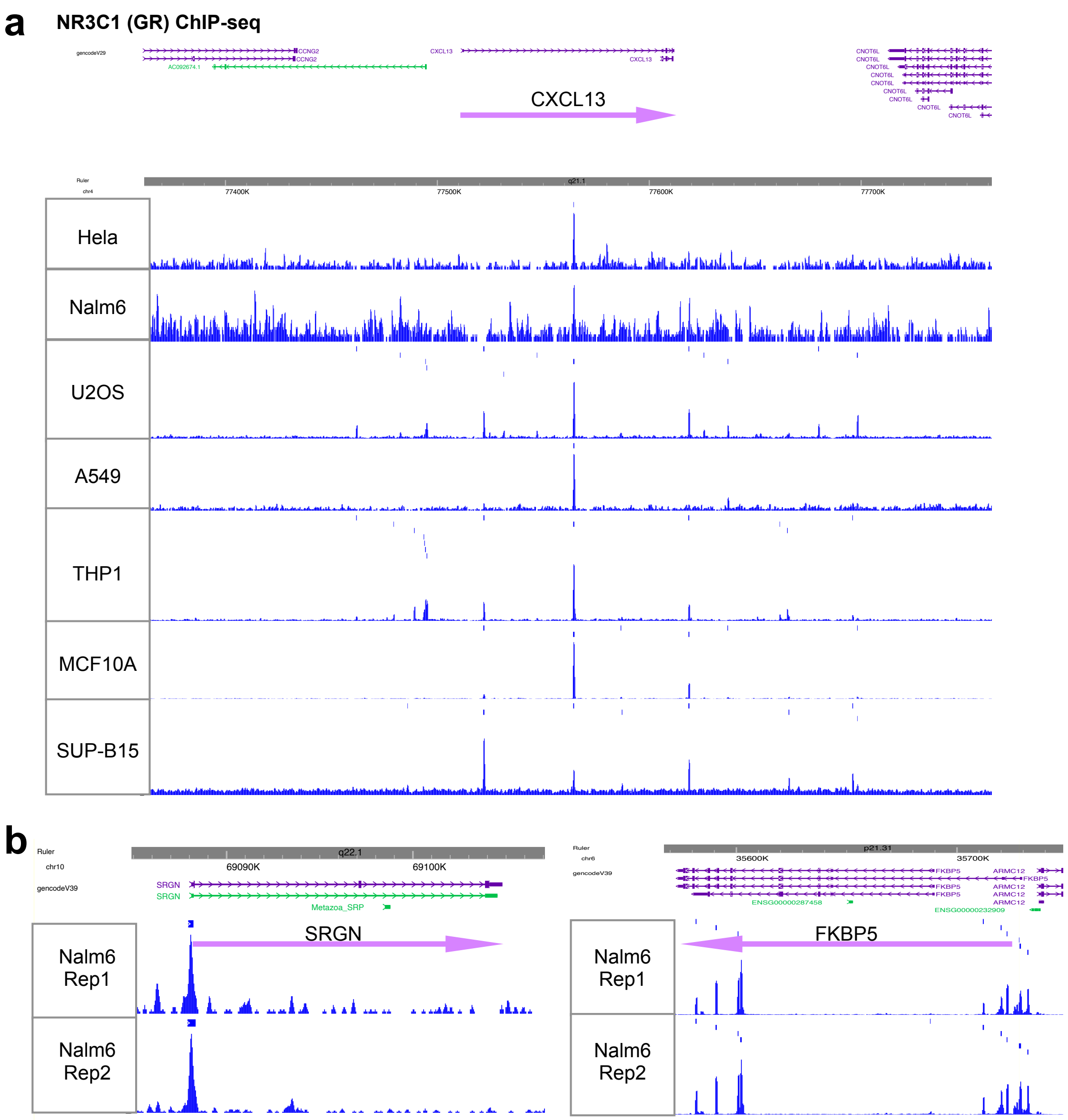


**Fig.S13. ChIP-seq binding of NR3C1. (a)** Glucocorticoid receptor (GR) ChIP-seq data from multiple cell lines including peaks and binding signals around CXCL13. **(b).**GR ChIP-seq data from Nalm6, a B cell leukemia line, binding at promoters of SRGN and FKBP5.

**Table S1. Meta-component interpretation. Meta-component Annotation:** Brief naming of each meta-component (MeC), with top 20, 100 genes by MeC z-weight. Annotation nonmeclature: "Category_CellState-Features". The first field before the underscore symbol marks 7 colored category of MeCs, 6 related to lineages: B for “B cell” , T for “CD4T, CD8T, NK cells”, DC for “Dendritic cell”, M for “Monocytes and macrophages”, Myeloid for “Other Myeloid types besides monocytes and macrophages”, Stroma for “Stromal cells", and the last category related to pan-celltype signaling pathway: Pan for "Pan-cell signaling pathway". **Meta-component Enrichment**: The significant enriched terms using various libararies using Enrichr and top 100 genes, as well as using GSEA and ranks of all genes. **Meta-component Regulator:** prioritized regulators based on either MeC z-weight or Lisa epigenetic siginificance. Red and Orange factors: TFs both highly weighted in MeC and significant in Lisa. Green: TFs significant only in MeC but not in Lisa binding. Blue: TFs significant in Lisa binding but not MeC z-weight.
